# Exploring viral particle, soil, and extraction buffer physicochemical characteristics and their impacts on extractable viral communities

**DOI:** 10.1101/2023.11.13.566929

**Authors:** Jane D. Fudyma, Anneliek M. ter Horst, Christian Santos-Medellín, Jess Sorensen, Grant Gogul, Luke Hillary, Sara Geonczy, Jennifer Pett-Ridge, Joanne B. Emerson

## Abstract

Soil viruses are expected to be pivotal members of soil ecosystems, and recent advances in viral size fraction metagenomic (viromic) approaches have substantially improved our ability to interrogate soil viral ecology. However, the first step of viromics relies on extraction buffers to effectively remove viral particles from the soil matrix for downstream analysis, and viral extraction efficiency at this stage could be affected by the interplay between viral particles, soils, and extraction buffer chemistry. Here, we investigated whether extraction buffer chemistry affected extractable viral community composition measured by viromics from different soil types, for both biological (samples collected 1 meter apart) and technical (subsamples from the same soil homogenate) replicates. We first investigated protein-supplemented phosphate-buffered saline pH (PPBS, pHs 4.5, 5.5, 6.5, and 7.5) on a forest, grassland and wetland soil that exhibited different soil edaphic properties, and then we tested different buffer chemistries (PPBS, Carbonated Buffer, Glycine, and Saline Magnesium) on just the wetland soil. Spatial distance, or where the sample was taken in the field (i.e., biological replicate), was the primary driver of extractable viral community composition across all buffers and soils tested. Differences in viral community composition according to extraction buffer properties were only observed in the grassland technical replicates at PPBS buffer pH 4.5, as well as in both the wetland technical and biological replicates treated with different buffer chemistries, but the effects of buffer chemistry were secondary to spatial distance in all cases where spatial distance was a factor. The lack of buffering capacity in the grassland soil technical replicates likely increased sorption of some viral particles at pH 4.5, but neither protein composition nor isoelectric point (both calculated bioinformatically) explained this phenomenon. Given that most soil viral ecological studies to date include sample collection schemes over distances much farther apart than the 1-meter distances considered here, results suggest that extraction buffer chemistry is likely of much lower importance than ecological considerations, such as spatial distance, in the design of future soil viral ecological investigations.

**HIGHLIGHTS:**

- Spatial distance was the main driver of extractable viral community composition.
- Extraction buffer chemistry secondarily structured wetland viral communities.
- At pH 4.5, PPBS buffer likely increased viral sorption in homogenized grassland soil.
- Increased sorption was not explained by estimated viral protein isoelectric points.

## 1. INTRODUCTION

Soil microorganisms are responsible for most nutrient transformations in soil, and viruses that infect these microorganisms presumably also exert controls on soil biogeochemical cycling processes. For example, in well-studied marine systems, viruses infect up to 10^23^ cells per second (Suttle, 2007), with substantial top-down controls on nutrient cycling and the structuring of microbial communities (Breitbart et al., 2018; Brum and Sullivan, 2015; Breitbart, 2012). Given that each gram of soil can harbor up to ten billion viral particles (Williamson et al., 2017), rivaling bacterial abundances (Torsvik and Øvreås, 2002), soil viruses are thought to have similar ecological impacts to their marine counter parts (Williamson et al., 2017; Kuzyakov and Mason-Jones, 2018). However, still very little is known about the composition of soil viral particles and communities or their functions in soil systems.

Our limited understanding of viruses in soil thus far is largely due to methodological challenges, which have recently been overcome. The first hurdle of moving marine viral ecology methods to a solid-centric biome (i.e., soil) was the difficulty of separating viral particles from the soil matrix for downstream analysis (Williamson et al., 2003; Emerson, 2019). Early viral ecology research employed various extraction buffers to mobilize viral particles from the soil, followed by 0.22 µm filtration to remove larger organisms, for both microscopic and genomic analyses (Ashelford et al., 2003; Williamson et al., 2003, 2005; Fierer et al., 2007; Kimura et al., 2008; Srinivasiah et al., 2013; Zablocki et al., 2014; Adriaenssens et al., 2015). While this viral particle extraction method is still the standard in soil viral ecological research today (Roux and Emerson, 2022), this approach was previously limited due to the state of library construction and sequencing technology in the recent past; the small amounts of recoverable viral DNA from viromics were insufficient for next-generation sequencing (Emerson, 2019). Thus, either various DNA amplification approaches were employed (Kim et al., 2008; Srinivasiah et al., 2013), or whole-community sequencing via first-generation sequencing was used (Fierer et al., 2007; Reavy et al., 2015), both which have substantial limitations, including biases towards easily amplifiable viruses (Yilmaz et al., 2010; Roux et al., 2016) and/or limited sequencing depth that precludes adequate recovery of viral community diversity (Roux et al., 2017). To circumnavigate these limitations, total metagenomes have recently been mined for viral sequences, greatly expanding our understanding of soil viral diversity (Emerson et al., 2018; Trubl et al., 2018; Khot et al., 2020). However, soils harbor substantial bacterial and extracellular DNA (i.e., genetic material from dead/inactive microbes, (Carini et al., 2016)), making viral genome recovery difficult; most soil metagenomes contain less than 1% recognizable viral content (Santos-Medellín et al., 2021; ter Horst et al., 2021). Advances in library construction (the preparatory steps for sequencing extracted DNA) have recently made it possible to sequence the small amounts (as little as a few ng) of recoverable DNA from viral-size fraction metagenomes (viromes), using next-generation sequencing approaches (Emerson, 2019; Trubl et al., 2020; Roux and Emerson, 2022). By employing viral particle separation and purification steps, with DNase treatment to remove extracellular DNA, viromics has emerged as one of the most effective methods to assess viral ecological patterns across diverse soil habitats (Trubl et al., 2019; Santos-Medellín et al., 2021; Roux and Emerson, 2022).

While viromics is an advantageous tool for viral community analyses, it hinges on the efficient recovery of diverse viral particles from the soil matrix. However, viral interactions with soil are dictated by both soil solution and soil surface chemistries (Jin and Flury, 2002), which can contribute to the efficiency of viral particle extraction by viromics. Soil pH, ionic concentration of the soil solution, organic matter content, and mineral composition can all drive the degree of viral sorption to soil surfaces (Gerba, 1984; Powelson and Gerba, 1995; Dowd et al., 1998; Jin and Flury, 2002; Williamson et al., 2003; Azadpour-Keeley and Ward, 2005), and the effects of these physicochemical differences can vary drastically across soil types (Voroney, 2007; Weil and Brady, 2019). The core assumption of viromics is that, by adding specific buffers that disrupt viral adsorption to soil particles, a representative sample of the viral community can be efficiently extracted from the soil matrix (Trubl et al., 2020). As such, researchers have been cautioned to tailor extraction buffers to different soil types, in order to extract a representative viral community, and biased viral particle extraction has been highlighted as a potentially serious limitation of the viromics approach (Trubl et al., 2020). Yet, the effects of buffer chemistry on extractable viral particles have been only disparately investigated under various soil-buffer combinations, typically linked to microscopic measurements of viral particle abundances and rarely to downstream sequencing data (Williamson et al., 2003, 2013; Trubl et al., 2016; Pratama and van Elsas, 2018; Göller et al., 2020), such that the field would benefit from more rigorous testing of the effects of different buffer chemistries on the diversity of recoverable viral communities from different soils.

In addition to the dynamic soil properties that control viral extraction efficacy, viruses themselves also have diverse physicochemical properties that contribute to their sorption potential, which can affect which viruses are extracted from soil by viromics. Viral surface proteins vary in their strength and direction of charge, based on the environmental pH (Michen and Graule, 2010), and it has been suggested that the isoelectric point (IEP), or the pH at which a virus is neutrally charged, can be a good indicator of viral desorption likelihood (Trubl et al., 2016, 2020). Yet, viruses have a large distribution of IEPs (Michen and Graule, 2010), and thus, different viruses may be more likely to be extracted at different buffer and/or soil pHs. Electrostatic forces are not the only factor driving viral sorption. Hydrophobic groups on the surface of viruses, competing environmental proteins, and viral particle size can also all contribute to viral extraction efficiency from soil (Powelson and Gerba, 1995; Jin and Flury, 2002; Ghernaout and Elboughdiri, 2021; Heffron and Mayer, 2021). Thus, a rigorous interrogation of the impacts of buffer and soil chemistry on recoverable viral communities, especially with buffers that differ in pH, ionic concentrations, protein composition, and hydrophobic-reducing agents, is necessary for appropriate interpretations of soil viromic datasets.

Here, we investigated whether extraction buffer chemistry affected the types and abundances of viruses extracted by viromics from different soil types. We first tested four different pHs of protein-supplemented phosphate-buffered saline solution (PPBS), a commonly used viromics extraction buffer (Göller et al., 2020; Santos-Medellín et al., 2021, 2022, 2023; Sorensen et al., 2021, 2023; ter Horst et al., 2021, 2023; Durham et al., 2022; Emerson et al., 2022), on forest, grassland, and wetland soils that exhibited distinctive physicochemical properties. We also bioinformatically explored the extracted viral particle surface protein composition and corresponding isoelectric points of these proteins under different extraction buffer pHs. We then tested the effects of four extraction buffers with different chemical compositions on a wetland soil. These buffers varied in pH and ionic composition, as well as whether amino acids and proteins were added to control hydrophobic interactions and site-specific competition with viral particles. For both sets of tests (comparisons according to PPBS buffer pH or buffer chemical composition), we analyzed both presumed biological replicates (soil cores collected approximately 1 m apart) and technical replicates from the same soil homogenate. These analyses of the effects of buffer chemistry on extractable viral particle compositional diversity were conducted to assess whether substantial methodological biases might underlie viromics-based soil viral ecological studies.

## 2. METHODS

### 2.1. Sample Collection

Samples were collected separately for each of four experiments (described in 2.2.), which, briefly, tested the impacts of buffer pH or buffer chemical composition on either biological or technical replicate samples. For all sets of biological replicates, samples were collected from the same soil habitat approximately 1 m apart, independently homogenized, and analyzed as three separate samples. For all sets of technical replicates, triplicate samples were collected from the same soil habitat approximately 1 m apart, homogenized together, and then separated into three (pH study) or four (buffer chemistry study) samples for analysis. For the first buffer pH experiment, samples were collected as biological replicates from each of three soil habitats in March of 2021, and the second pH experiment used technical replicates collected from the same habitats in December of 2021. The sampled sites were a mixed conifer forest (‘forest’) located at the Angelo Coast Range Reserve (39.727995, -123.645758), a Mediterranean grassland (‘grassland’) located at the McLaughlin Natural Reserve (38.875519, -122.418847), and a freshwater wetland (‘wetland’) located at the Bodega Marine Reserve (38.318898, - 123.069616). For the two buffer chemistry experiments, samples were collected at the same freshwater wetland as for the pH study, first in November 2021 (technical replicates, n= 4) and second in November 2022 (biological replicates, n=3). All samples were collected as surface cores (0-15 cm deep, 2.5 cm diameter) and processed within four days of sampling.

### 2.2. Experimental Setup and Buffer Preparation

For the buffer pH experiments, protein-supplemented phosphate-buffered saline (PPBS) buffer was prepared at four different pHs, each of which was used for viromics on biological (first pH experiment) and technical (second pH experiment) replicates from three soil types (forest, grassland, wetland). For each soil, each biological replicate (n=3) was homogenized, and then subsamples of each homogenate were treated with each of the four buffer pHs. Three technical replicates per soil were treated with each of the four buffer pHs. PPBS was prepared as previously published (Göller et al., 2020; Emerson et al., 2022), and was composed of 2% bovine serum albumin, 10% phosphate-buffered saline, 1% potassium citrate, and 150 mM MgSO_w_. The pH of each PPBS buffer was adjusted with either sterile 2N HCl or 6M K_2_CO_3_, for final pH concentrations of 4.5, 5.5, 6.5 (control), and 7.5. For the buffer chemistry comparison experiments, biological and technical replicates of the wetland soil were tested under four different extraction buffers. The four buffers were adapted from previous protocols and prepared to the following concentrations: Carbonated Buffer (CB, 60 mM NaHCO_3_, 100 mM NaCl, 3 mM KCl, final pH 8.65 (Zhuang and Jin, 2003)), Glycine (GL, 250 mM, final pH 6.34 (Williamson et al., 2003)), and Saline Magnesium (SM, 100 mM NaCl, 8 mM MgSO_4_, 50 mM Tris (pH 7.5), final pH 7.83 (Thurber et al., 2009)). All solutions were sterilized via autoclave and 0.22 µm filtration. PPBS adjusted to pH 6.5 was used as the fourth (control) buffer for the buffer chemistry comparison experiments. Subsamples of each homogenized biological replicate were treated with each of the four buffers (n=12), and each of four technical replicates was treated with the four distinct buffers (n=16).

### 2.3. Virome DNA extraction, library construction, and sequencing

Soil virions were purified and extracted as previously described (Emerson et al., 2022; Santos-Medellín et al., 2022), apart from the differences in buffer pH or buffer chemical composition described above. Briefly, for each sample, 10 g of soil were suspended in 9 mL of buffer, briefly vortexed, agitated for 10 min on an orbital shaker (300 rpm at 4°C), centrifuged (10 min, 4000 RPM, 4°C), and supernatant was poured into a new tube. This was repeated on the same soil twice more, for a total volume of 27 mL of supernatant. Supernatant was centrifuged twice to remove residual soil particles (8 min, 7840 RPM, 4°C) and then filtered through a 0.22 µm filter to remove most cells. The filtrate was ultracentrifuged under vacuum for 2 hours and 25 minutes (35,000 RPM, 4°C, 50.2 Ti rotor) to pellet the virions. Pellets were resuspended in 100 µL of UltraPure water and treated with Promega RQ1 RNase-Free DNase (1:10 ratio DNase to solution, 30 minutes, 37°C). Viral DNA was extracted from the DNase-treated virions using the DNeasy PowerSoil Pro kit (Qiagen) according to the manufacturer’s instructions, with an additional heat treatment (10 min, 65°C) before the bead-beating step. Libraries were constructed by the University of California, Davis DNA Technologies Core, using the DNA KAPA HyperPrep library kit (Roche). Paired-end 150 bp sequencing was generated using the NovaSeq S4 platform (Illumina) to a depth of at least 10 Gbp per virome.

### 2.4. Soil Physicochemical Analyses

All soil physicochemical analyses were performed by Ward Laboratories (Kearney, NE, USA), as follows: soil pH and soluble salts (1:1 soil water ratio), total organic matter (loss on ignition %), nitrate-N (KCl extraction), phosphorus (Olsen P), potassium, calcium, magnesium, and sodium (ammonium acetate extraction), sum of cations (CEC, calculated from K, Ca, Mg, Na and buffer pH), cation base saturation (calculated from CEC), sulfate-S (Mehlich 3 extract), copper, iron, manganese, and zinc (DTPA extraction) (**Table S1**).

### 2.5. Soil pH Changes

The pHs of the buffers and soils were measured at three time points during the buffer pH technical replicate experiment. Buffer pH and soil pH were measured separately before the experiment was conducted. The pH of the buffer was measured directly, while soil pH was measured using a 1:1 soil-water ratio. After the addition of the 27 mL of buffer and shaking steps, the pH was measured of the buffer supernatant before it underwent any further centrifugation steps. Finally, the pH of the remaining soil that had undergone extraction was measured, after supernatant removal, again using a 1:1 soil-water ratio.

### 2.6. Virome Assembly, Identification and Quantification

Raw virome reads were quality filtered using Trimmomatic v0.39 (Bolger et al., 2014), with a minimum quality-per-base score of 30 and a minimum read length of 50 bases. PhiX sequences used in the sequencing process were removed using BBduk from the BBMap package (Bushnell, 2014). *De novo* assemblies were generated for each virome separately, using MEGAHIT v1.0.6 (Li et al., 2015) in meta-large mode and requiring 10,000 bp minimum contig length. Contigs were predicted as viral using VIBRANT v1.2.1(Kieft et al., 2020) with the virome flag. Viral species-level dereplication (i.e., generation of the set of vOTUs) was performed using dRep v3.2.2 (Olm et al., 2017) on only the medium- and high-quality viral contigs identified by VIBRANT, where viral contigs were dereplicated at 95% average nucleotide identity (ANI) with a minimum coverage threshold of 85%, using the ANImf and single cluster algorithms. Reads were mapped against the database of dereplicated vOTUs in sensitive mode, using Bowtie2 v2.4.2 (Langmead and Salzberg, 2012) and, due to differences in sequencing depth, each virome was rarefied by randomly subsampling the mapped reads by a factor based on the lowest sequencing depth, using the view function in SAMtools v1.14 (Li et al., 2009). A mean coverage table of both the rarefied and non-rarefied data was generated using CoverM (Woodcroft, 2023) with a minimum contig breadth coverage threshold of 75% and a minimum read percentage identity of 90%.

### 2.7. Viral Gene-sharing Taxonomy Network

Protein content for each vOTU recovered in the biological and technical replicates from the pH experiment was predicted using Prodigal v2.6.3 (Hyatt et al., 2010). The output amino acid file was used to build a gene-sharing network using vConTACT2 v0.9.19 (Jang et al., 2019). Briefly, Diamond (Buchfink et al., 2021) was used to perform the protein alignment, and the MCL algorithm was used to cluster proteins. The clustered vOTUs were visualized using Cytoscape v3.9.1 (Shannon et al., 2003), with vOTUs colored either by the soil type or buffer pH treatment from which they were assembled, or by significant differential abundances across pHs (see section 2.9.).

### 2.8. Viral Surface Protein Prediction and Isoelectric Point Calculations

A viral surface protein database of amino acids was curated from the vConTACT2 viral RefSeq prokaryotes protein database (version 211, April 2022 (Jang et al., 2019)). The vConTACT2 amino acid sequences were filtered based on keywords associated with viral surface proteins (i.e., capsid, membrane-associated, tail, spike) and manually inspected to ensure that selected sequences were consistent with viral surface moeities or structural elements. Briefly, proteins with annotations containing terms such as ‘assembly factor’, ‘protein precursor’, ‘assembly protein’, and ‘inner membrane protein’ were removed from the database, whereas terms such as ‘major capsid protein’, ‘integral membrane protein’, ‘coat protein’, and ‘outer capsid protein’ were retained (**Table S2**). Viromic sequencing reads that had mapped to vOTUs were then aligned against the surface protein database amino acid sequences, using PALADIN (Westbrook et al., 2017). The resulting bam files were filtered to only include entries with the highest maximum mapping quality score, as recommended by the program. A mean surface protein coverage table was produced from the mapping files, using CoverM (Woodcroft, 2023) with the same settings used for generating the vOTU abundance table. Theoretical isoelectric points of amino acid sequences were calculated, using the computePI function in the seqinr package (Charif and Lobry, 2007). Briefly, the computPI function calculates isoelectric points from pK values of all amino acids provided in a protein sequence (Bjellqvist et al., 1993).

### 2.9. Data Analyses and Visualization

All data analyses were performed in R (version 4.2.3) on the ≥ 75% breadth coverage, mean (75-mean) coverage vOTU table, and most data wrangling was performed using Tidyverse v2.0.0 (Wickham and RStudio, 2023). Vegan v2.6.5 (Oksanen et al., 2022) was used to calculate Bray-Curtis dissimilarities on Hellinger transformed relative abundances, and the dissimilarity matrix was used for Principal Coordinates Analyses (PCoA), using the Ape package v5.7.1 (Paradis and Schliep, 2019). PERMANOVA was performed on dissimilarity matrices, using the adonis2 function in Vegan (**Table S3**), followed by the pairwiseAdonis v0.4.1 (Arbizu, 2020) post-hoc test. Viral richness was calculated on the rarefied vOTU 75-mean abundance table, using the Vegan package, and checked for normality of distribution using the Shapiro-Wilk’s test (base R), and density- and qq-plots (ggplot2 v3.4.3, (Wickham et al., 2023)). To test for significant differences in richness, either one of two methods were followed. Normal data was fitted using a linear model (base R), followed by ANOVA (base R, **Table S4**) and Tukey pairwise post-hoc tests (emmeans package v1.8.9 (Lenth et al., 2023)). Non-normal data was fitted using a generalized linear model (glm) with “poisson” family distribution (for positive count data, base R), followed by an analysis of deviance (ANOVA with “Chisq” test, base R, **Table S5**) and Tukey pairwise post-hoc tests (emmeans). This same statistical approach (checking for normality, model fitting, testing significance of predictor variables, post-hoc testing) was used on the DNA yield and isoelectric point data, except non-normal, glm fitted models specified “gamma” distribution as family type (for continuous positive integer distributions, **Table S5**).

Differential abundance analysis was performed across all pairwise pH comparisons (i.e., 4.5 vs 5.5, n = 7) on vOTU counts using DESeq2 v. 1.38.3 (Love et al., 2014), and the pairwise Wald test p-values were manually adjusted to account for all pairwise comparisons using the Bonferroni method. Distance-decay relationships were plotted, using linear regressions between the surface protein composition Bray-Curtis dissimilarity and absolute distances between pHs in pairs of samples, and goodness of fit was determined using Pearson’s correlations. Venn diagrams (ggvenn v0.1.10 (Yan, 2023)) were calculated from presence-absence transformed, rarefied, 75-mean coverage tables, where only vOTUs present in all samples for a given group (i.e., all replicates, all buffer pHs, or all buffer types) were used for analysis. Soil physicochemical data was z-score transformed, and the Euclidian distances were visualized, using Principal Components Analysis (Vegan). Pheatmap v1.0.12 (Kolde, 2019) was used to cluster and visualize the z-score transformed soil physicochemical data, using the complete clustering method and default settings. Statistical differences in soil physicochemical properties between two sampling dates were checked for normality (as stated previously), followed by either an unpaired T-test (normal data) or Wilcoxon rank sum test (non-normal data, base R, **Table S6**). All plots were generated with ggplot2 v3.4.3, ComplexUpset v1.3.3, ggvenn, and cowplot v1.1.1 (Wilke, 2020; Krassowski, 2021; Wickham et al., 2023; Yan, 2023).

## 3. RESULTS

### 3.1. Considering presumed biological replicates, recovered soil viral communities were significantly structured by both soil type and spatial distance but not by PPBS extraction buffer pH

To determine whether extraction buffer pH substantially controlled the types and relative abundances of viruses extracted and sequenced from soil, we tested four pHs (4.5, 5.5, 6.5, and 7.5) of PPBS extraction buffer on three soils (a forest, a grassland, and a wetland), first considering presumed biological replicates (near-surface soils collected approximately 1 m apart within each site). The three different soil sites were chosen to capture a range of soil physicochemical properties (**Figure S1A**), which might affect both the chemistries of viral particles adapted to each ecosystem and the interactions of the soil and its viral community with our extraction buffer across the tested pH range. The specific buffer pHs were chosen because they are within the range of pHs observed naturally in the three soil systems. Putative biological replicates were used for this first experiment, as we hoped that they would capture similar viral community composition, as well as account for some microheterogeneity within each soil site. We recovered 8,490 viral operational taxonomic units (vOTUs) across soils and experimental units (different buffer pH and biological replicate combinations). Considering that the tested soils had a wide range of soil edaphic properties (e.g., soil organic matter content, cation composition, and macronutrients **Figure S1B, Table S1**), it was unsurprising that viral communities differed most significantly across the three soil habitats, regardless of extraction buffer pH used (PERMANOVA p = 0.001, **Figure 1A, Table S3**). Further, when assessing the total number of vOTUs recovered in a single soil type, more than 90% of the vOTUs were only detected within that soil type, with zero vOTUs shared across all three soil sites (**Figure 1B**).

**Figure 1.**
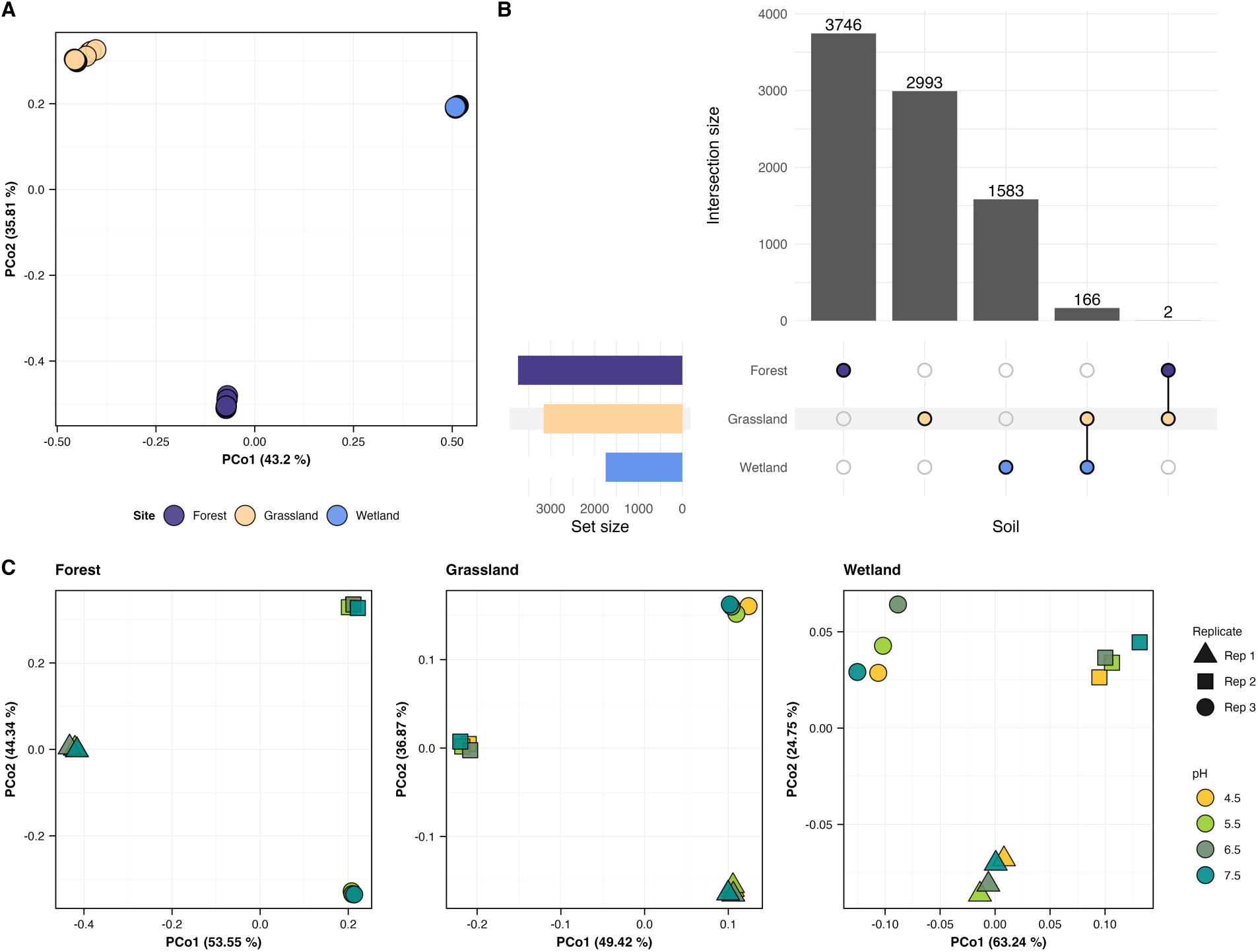
Extractable viral community composition patterns for biological replicates treated with different PPBS buffer pHs. **A)** Principal coordinates analysis (PCoA) based on Bray-Curtis dissimilarities of vOTU abundances (coverage depths from read mapping) from three soils, with 12 viromes per soil (three biological replicate samples collected ∼1 meter apart, each treated with protein-supplemented phosphate-buffered saline solution (PPBS) at four different pHs: 4.5, 5.5, 6.5, and 7.5). Each point is one virome, representing one biological replicate and pH treatment combination, colored by soil type. **B)** UpSet plot of shared vOTUs in each soil type, based on presence-absence data derived from read mapping. Colored dots indicate the soil type(s) in which a given set of vOTUs was detected, and connecting lines between dots indicate vOTUs shared between soils. Data for each soil is derived from 12 samples (three biological replicates and four PPBS buffer pHs), where any vOTU detected in any sample was counted for this analysis. **C)** Principal coordinates analyses (PCoA) based on Bray-Curtis dissimilarities of vOTU abundances (coverage depths from read mapping), faceted by soil type. Each point is one virome. For each soil type, each virome is labeled according to biological replicate (shapes, Rep 1, Rep 2, Rep 3) and colored by PPBS buffer pH.

The forest soil had the highest vOTU richness, followed by the grassland and then the wetland soils (**Figure 1B**). Although species-level viral communities and total diversity were significantly different by soil type, genus-level viral clusters did not reflect any differences according to soil type or extraction buffer pH (**Figure S2A,B**). These results together confirm that our experiments captured very different edaphic properties and species-level viral communities in each of the three soil habitats, suggesting that we might expect the impacts of different buffer pHs on extractable viral communities to differ across sites, or, if not, that the results might be broadly generalizable to other soil types.

Despite our expectations, when comparing within each soil type, viral community composition was most significantly different among presumed biological replicates taken ∼1 m apart for all soils tested (PERMANOVA p < 0.05, **Table S3**), while extraction buffer pH had no significant effect on extracted viral community composition (**Figure 1C**). Similarly, viral richness was significantly different between biological replicates (BR-1 and BR-2) in both the grassland and wetland soils (**Figure S3B**), and viral particle abundances (measured by proxy as total viromic DNA yields (Santos-Medellín et al., 2023)) were significantly higher in one out of the three grassland soil biological replicates (**Figure S4B**). The only observed effects of PPBS extraction buffer pH was in the wetland soil, where viral richness was significantly lower in pH 6.5 as compared to pH 4.5 (**Figure S3A**). Taken together, these results show that extraction buffer pH did not significantly affect viral community composition or viral particle abundances in any of the soils, and viral richness differed at only one buffer pH in one soil type. The significant differences in viral community composition among presumed biological replicates in all three soils, as well as significant differences in viral richness between presumed biological replicates in the grassland and wetland soils, indicate that a 1 m distance between soil samples is not sufficiently short to capture substantially overlapping viral communities, such that these were not biological replicates after all. Though these differences in viral community composition over relatively short spatial distances were discovered here accidentally, this experiment demonstrates that the ability to detect these local biogeographical patterns was not affected by extraction buffer pH in any of the tested soils.

### 3.2. Recovered soil viral communities from technical replicates did not differ significantly across PPBS buffer pHs in forest or wetland soils but did differ significantly at the lowest buffer pH in the grassland soil

As the strong spatial structuring of viral community composition could have masked the effects of buffer pH on extractable viral community composition in the experiment with the presumed biological replicates, we next sought to control for spatial heterogeneity by using technical replicates. We tested the same four PPBS buffer pHs on three subsamples of the same soil homogenate from each site. Sampling was the same as for the ‘biological replicates’, with three near-surface soil cores collected approximately 1 m apart, but those triplicate cores were homogenized for each site to generate three technical replicates per site. In this experiment, we recovered 6,569 vOTUs in total. As expected, and consistent with the biological replicate experiment, viral communities differed most significantly by soil type (**Figure S5A**), and zero vOTUs were shared across all three soils (**Figure S5B**). When considering all three soils together, extraction buffer pH did not have a significant effect on extracted viral community composition (PERMANOVA p = 0.206, **Table S3**), consistent with the substantial differences in viral community composition among the three soils.

Some effects of extraction buffer pH on extractable viral community composition, richness, and/or abundance across technical replicates became apparent within soil types, but these effects were different for each soil. For the forest and wetland soils, viral community composition did not differ significantly by extraction buffer pH, but grassland viral communities extracted at pH 4.5 were significantly different from the communities extracted at pHs 5.5 and higher (PERMANOVA p = 0.017, **Figure 2A**, **Table S3**). Further, extracted viral richness at pH 4.5 in the grassland technical replicates was significantly lower than at pHs 6.5 and 7.5 (**Figure 2B**), with a similar, significant trend observed in the overall abundance of extracted viral particles (lowest at pH 4.5), using viromic DNA yields as a proxy for viral abundance (**Figure S4C**). The forest and wetland soils also showed decreases in viromic DNA yields (extractable viral particle abundances) with decreasing buffer pH (**Figure S4C**), but viral richness in those two soils did not significantly differ across extraction buffer pHs (**Figure S3C**). Given the lower viral richness and DNA yields at pH 4.5 in the grassland soil, we also investigated whether the relative abundances of vOTUs were significantly lower between pH 4.5 and the other pH groups. The grassland soil had the highest number and proportion of significantly differentially abundant vOTUs between pH treatment groups (i.e., 4.5 vs 5.5-7.5), where 155 vOTUs were depleted in pH 4.5, and only 32 vOTUs were enriched, comprising 6.9% and 1.4% of the recovered vOTUs in the grassland soil, respectively (**Figure 2C**). The forest soil followed a similar trend, but for a much smaller subset of vOTUs (2.6% of all recovered vOTUs), and the wetland soil showed no obvious pattern between pH groups (**Figure S6**). Considering ‘genus-level’ viral clusters (Jang et al., 2019), there was no clustering of vOTUs based on depletion or enrichment in pH 4.5 for the grassland soil (**Figure S2C**). Altogether, only the grassland soil extracted with a buffer pH of 4.5 showed significant differences in extractable viral community composition, richness, and abundances; all other buffer pHs for the grassland and all four buffer pHs for the other two soils had no significant effect on any of the measured viral extraction parameters. Still, most grassland vOTUs (81.2%) were extracted at all four buffer pHs, with less than 1% uniquely extracted at pH 4.5 (**Figure S7B**).

**Figure 2.**
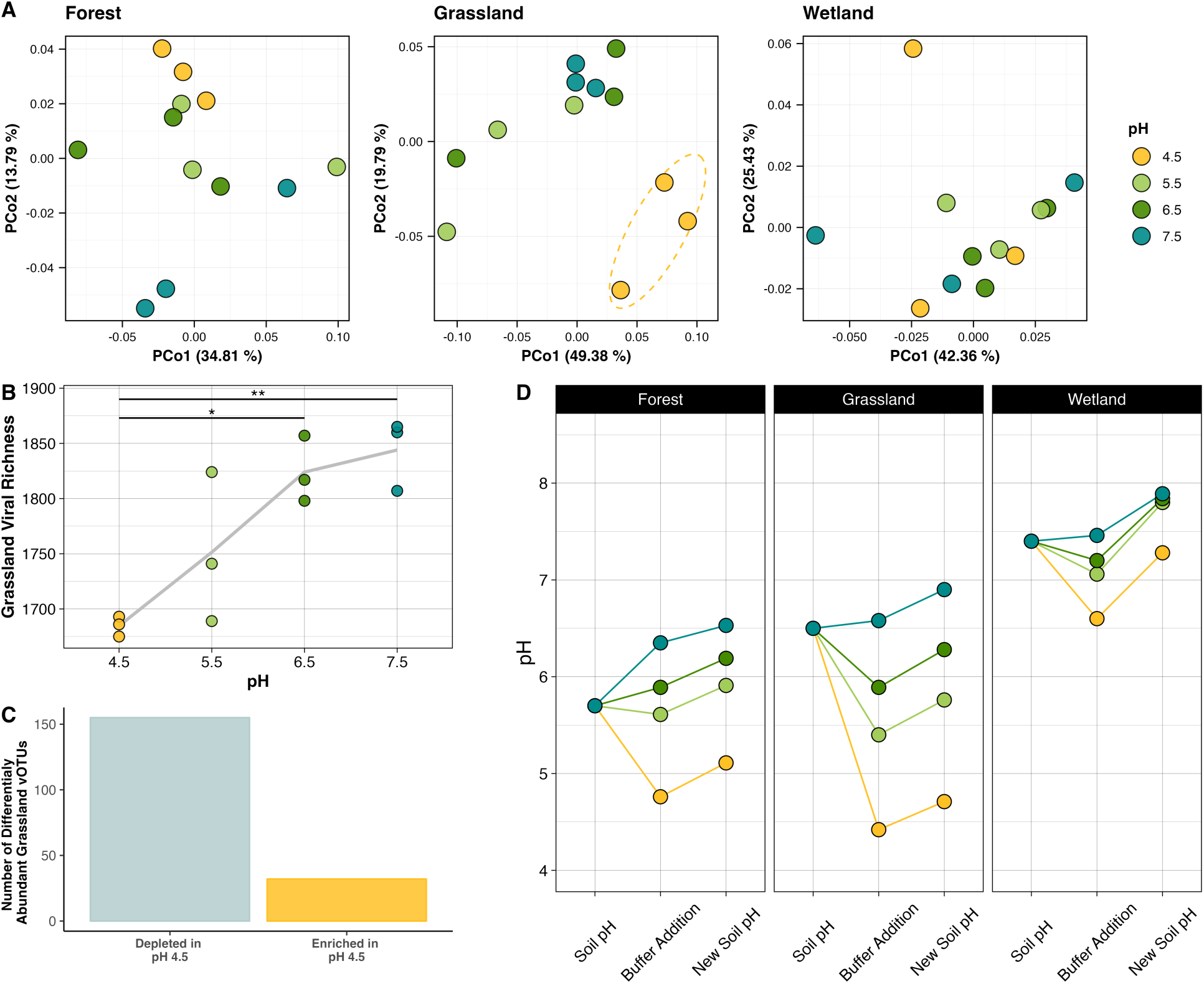
Extractable viral community composition patterns and soil responses for technical replicates treated with different PPBS buffer pHs. **A)** Principal coordinates analysis (PCoA) based on Bray-Curtis dissimilarities of vOTU abundances (coverage depths from read mapping) from technical replicates (samples derived from single within-site homogenates) treated with protein-supplemented phosphate-buffered saline solution (PPBS) at four different pHs, faceted by soil type. Each point is a virome from one technical replicate and pH treatment combination, colored by PPBS buffer pH. Dotted yellow line in the grassland PCoA indicates the 95% confidence interval ellipse for the pH 4.5 group. **B)** Viral community richness measurements (total numbers of vOTUs) from the grassland soil across PPBS extraction buffer pHs. Points represent individual viromes from one technical replicate and buffer pH combination, and the trend line follows the mean viral richness across buffer pHs. Significance between groups is marked by connecting lines with significance stars (*, p <0.05; ** p<0.01), based on analysis of variance (ANOVA) and Tukey post hoc test. **C)** Significantly differentially abundant vOTUs in the grassland soil technical replicates between the pH 4.5 extraction and all other pH extractions (5.5, 6.5, 7.5), reflecting vOTUs either enriched or depleted in the pH 4.5 extraction. **D)** Measured pH of: the soil before virome extraction, the supernatant after three extraction steps (containing buffer and extracted viral particles after buffer addition), and the remaining soil after the supernatant was removed for downstream viral particle extraction. Colors represent the pH of the PPBS buffer used.

We hypothesized that differences in soil buffering capacity (the ability to buffer against changes in pH (Weil and Brady, 2019)) might explain why differences in extractable viral community composition were only observed in the grassland soil at buffer pH 4.5. Thus, we measured the pH of the buffer supernatant containing viral particles after extraction from soil, which would reflect the immediate exchange of compounds and reveal whether the intended buffer pH was maintained throughout the viral particle extraction steps. We also measured the pH of the remaining soil after the extraction buffer was removed, which would reflect the residual acidity and, in comparison to the initial soil pH, the buffering capacity of the soil (i.e., soils with higher buffering capacity should remain close to the initial soil pH measured prior to buffer addition). Only the grassland pH 4.5 supernatant remained at pH 4.5 during the extraction steps, whereas the wetland pH 4.5 supernatant increased to pH 6.5 (**Figure 2D**). The forest supernatant increase to pH 4.75, which was close to the intended extraction pH, yet this soil started at a lower pH than the grassland soil (5.75 vs. 6.5), meaning that the forest soil should have reached the intended pH of 4.5 if the two soils had had similar buffering capacities.

Similarly, the final soil pH of the wetland after extraction was only ± 0.5 pH units from its original pH, and the forest soil showed a slightly larger pH change of 1 pH unit. The grassland soil, however, demonstrated the largest pH change, decreasing by nearly 2 pH units after the addition of pH 4.5 buffer. These results suggest that the wetland and forest soils were more capable of buffering against the addition of acidic buffer pHs, which is consistent with the soil edaphic properties exhibited in these two soils. More specifically, cation exchange capacity (CEC, i.e., the amount of cation sorption sites), which is largely influenced by organic matter content, is central to a soil’s ability to buffer against pH changes (Nelson et al., 2010; Curtin et al., 2013), and both the wetland and forest soils had 3.4- and 2.1-times greater CEC, as well as 4.25- and 8.9-times greater soil organic matter content (**Table S1**). It is therefore likely that differences in extracted viral communities across buffer pHs were only observed in the grassland soil due to a relative lack of buffering capacity in that soil, which facilitated larger shifts in pH after buffer addition, presumably resulting in a stronger influence of extraction buffer pH on viral sorption dynamics.

We wanted to further investigate the influence of pH on viral sorption dynamics by considering viral surface protein composition. For example, protonation of surface protein functional groups with increasing hydrogen ion concentrations (lower pHs) could accelerate sorption of certain viral particles, based on their surface protein makeup (Michen and Graule, 2010). This could result in different pools of viral particles with distinct protein compositions extracted over a pH range. We hypothesized that this should largely be observed only in the grassland soil, due to its lack of buffering capacity, which we presume was the underlying driver of the distinct composition, reduced diversity, and large number of depleted vOTU abundances recovered at pH 4.5. To estimate viral surface protein compositional differences across soils, we bioinformatically aligned viral reads to a database of viral surface protein amino acid sequences (**Table S2**), giving us the relative abundance of each surface protein in each sample, which we used to generate a pairwise distance matrix of predicted surface protein compositions.

Predicted surface protein compositions were significantly different among the three soils tested (PERMANOVA p = 0.001, **Figure 3A**, **Table S3**), yet, within each soil, the recovered surface protein composition was not significantly structured by PPBS buffer pH for any of the soils (**Table S3**). However, while not significant, the grassland soil exhibited the strongest correlation (steepest slope, highest R^2^ value, and lowest p-value) between protein compositional dissimilarity and distance between pH measurements (**Figure 3B**). This suggests that, in the grassland, extraction buffer pH selected for more distinct protein compositions across a pH range, potentially from different pools of viral particles more readily desorbed at different pHs.

**Figure 3.**
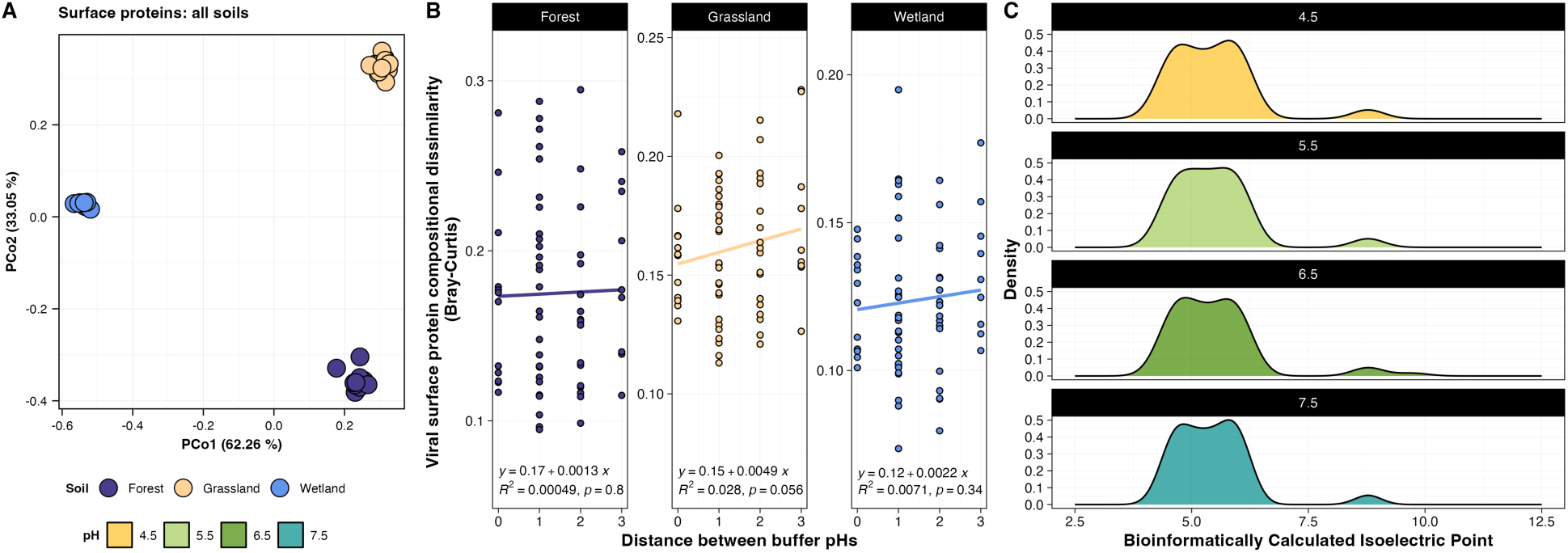
Surface protein compositional and predicted isoelectric point differences from vOTUs recovered in technical replicates treated with different PPBS extraction buffer pHs. **A)** Principal coordinates analysis (PCoA), based on Bray-Curtis dissimilarities of viral surface protein compositions in each virome from the ‘technical replicates’ dataset, derived from read alignments to predicted surface protein sequences. Each point represents surface proteins from one virome, colored by soil. **B)** Distance-decay trends of pairwise viral surface protein compositional dissimilarities by pairwise differences between PPBS buffer pHs for each soil type. Points represent pairs of samples and points are fitted with linear regression trend lines. The equation, linear regression p-value, and goodness of fit (R^2^) value for each trend line is reflected at the bottom of each graph. **C)** Density plot reflecting the distributions of bioinformatically calculated isoelectric points (i.e., the pH at which a protein is neutrally charged) from predicted viral surface proteins of vOTUs recovered from the grassland soil technical replicates, faceted by pH.

We also calculated the theoretical isoelectric points (IEPs) of the detected surface proteins to better understand charge-sorption potential across pHs. Specifically, the isoelectric point indicates the pH at which a protein is neutrally charged, such that these surface proteins would be expected to sorb to soil at pHs below their isoelectric points (Jin and Flury, 2002). As viral diversity and extractable particle abundances (using viromic DNA yields as a proxy) decreased with decreasing pH in the grassland soil, we expected that the IEPs would also decrease, reflecting the pool of desorbed viral particles with IEPs below the extraction pH. While the theoretical IEPs of the surface proteins in each soil were distributed differently (**Figure S8A**), the grassland soil IEP distributions were not distinct to each buffer pH, nor did they trend lower with decreasing pH (**Figure 3C**). There was also no significant difference in IEPs across pH in the other two soils (**Figure S8B, Table S5**). Taken together with the protein compositional data above, these results suggest that the differences observed only in the grassland viral communities extracted at pH 4.5 could be partially explained by the correlation between extraction buffer pH and surface protein composition, but the predicted charges of these proteins did not change over a pH range and thus do not readily support the enhanced sorption hypothesis.

### 3.3 Considering technical replicates from the wetland soil, viral community composition differed significantly by extraction buffer chemistry, but most vOTUs were extracted by all four buffers

Since pH had only slight effects on extractable viral abundances and community composition, we next decided to test whether buffers with completely different chemical compositions would affect extractable viral communities. We chose the wetland soil for this experiment because it contained the largest shared pool of viral species among ‘biological replicates’ (**Figure S7C**), or in other words, this soil demonstrated the smallest degree of spatial heterogeneity of the three soils tested. Thus, we believed that, in the wetland soil, extraction buffer chemistry may be more likely structure viral community composition, as spatial heterogeneity would not be a prominent confounding variable. Still, due to the strong spatial structuring observed in the first experiment, we first used technical replicates to test the effects of four different extraction buffers (carbonated buffer, glycine, saline magnesium, and PPBS at pH 6.5) on extractable viral community composition. This time, extractable viral community composition differed significantly by extraction buffer used (PERMANOVA p = 0.999 E-3, **Figure 4A, Table S3**). However, extracted viral richness was not significantly different across the four buffers (**Figure S9A**). Of the total extracted vOTUs, over 85% were extracted by all four buffers, and while PPBS extracted the most unique vOTUs, this was only by a marginal amount (1.5% of the total vOTUs extracted, **Figure 4D**). Viral particle abundances by viromic DNA yield proxy were slightly higher in the PPBS extraction, as well as in the saline magnesium extraction, but again, these differences were not significant (**Figure S9D**). Taken together, across technical replicates from a wetland soil, viral community composition, but not richness or particle abundances, differed significantly, according to the buffer chemistry used for extraction.

**Figure 4.**
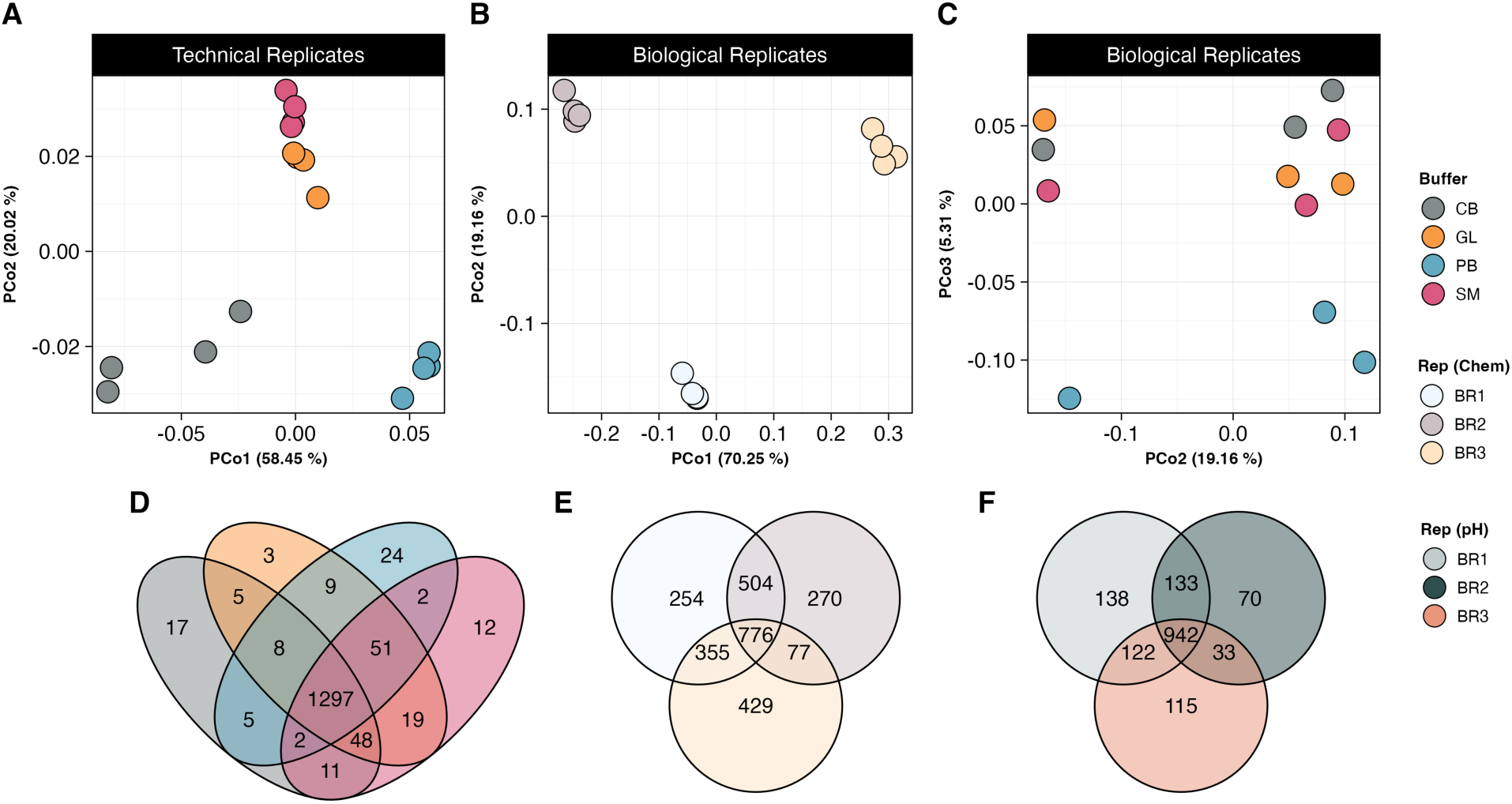
Extractable viral community composition patterns in a wetland soil treated with different extraction buffer chemistries. **A-C**) Principal coordinates analyses (PCoA plots) of viral community Bray-Curtis dissimilarities (from read mapping-derived vOTU abundances) in the wetland soil treated with four different buffer chemistries [CB – carbonated buffer, GL – glycine, PB – protein-enhanced phosphate buffered saline (PPBS, as in earlier tests, here pH 6.5), SM – saline magnesium]. Each point is one virome, where **A)** depicts technical replicates, colored by buffer chemistry (16 viromes from the same soil homogenate, 4 replicates per buffer), **B)** shows biological replicates (three samples, BR1-3, collected ∼1m apart) that each received all four pH chemistries, colored by replicate, and **C)** shows the second and third PCoA axes of the analysis from panel B (biological replicates), here colored by buffer chemistry. **D-F)** Venn diagrams of vOTU detection patterns (based on presence-absence data derived from read mapping to vOTUs) in the wetland soil from both the buffer chemistry and PPBS buffer pH experiments; vOTUs were only considered for this analysis if they were detected in all technical replicates for a given buffer treatment (D) or all treatments for a given biological replicate (E, F). **D)** vOTUs detected within or across different buffer chemistries for technical replicates from the buffer chemistry experiment, **E)** vOTUs detected within or across replicates for biological replicates from the buffer chemistry experiment, and **F)** vOTUs detected within or across replicates for biological replicates from the buffer pH experiment.

### 3.4. Considering biological replicates from the wetland soil, viral community composition differed most significantly among replicates and secondarily by extraction buffer chemistry

Given that buffer chemistry significantly altered wetland viral community composition across technical replicates, while spatial heterogeneity was the only significant driver of differences in wetland viral community composition across PPBS buffer pHs, we wanted to test whether the effects of buffer chemistry were preserved while introducing spatial heterogeneity as another variable. Thus, we tested the same four buffers on biological replicates (samples collected ∼1 m apart). In this case, biological replicate (i.e., where the sample was collected in the field) was the most significant driver of viral community composition, as observed across Axes 1 and 2 of a PCoA plot (PERMANOVA p = 0.001, **Figure 4B, Table S3**). However, the PERMANOVA analysis further revealed that buffer chemistry was also a significant driver of viral community composition (p = 0.024, **Table S3**), as can be visualized across Axes 2 and 3 of the PCoA analysis (**Figure 4C**). Additionally, viral richness and viral particle abundances (by DNA yield proxy) trended slightly higher in the PPBS buffer group relative to the three other buffers (**Figure S9C, S9F**), and PPBS extractions also recovered the most unique vOTUs of the four buffers tested (7.7% of total vOTUs, **Figure S9I**). However, despite the significant impact of buffer chemistry on wetland extractable viral communities in biological replicates, still more than half (58%) of the total extracted viral species were extracted by all four buffers, while less than a third (29.1%) of the total extracted viral species were shared across all three biological replicates (**Figure S9H,S9I**). Together, these results suggest that spatial distance and not buffer chemistry was still the primary driver of differences in extracted viral community composition.

Since buffer chemical composition influenced the extracted viral community composition among biological replicates in the wetland, but PPBS buffer pH did not, we hypothesized that the wetland buffer chemistry biological replicates might have more closely resembled technical replicates. In other words, we thought that the influence of buffer chemistry on biological replicates might only have been apparent because the viral communities were less spatially structured (better mixed) at the collection time point for the buffer chemistry experiment than they were when biological replicates were collected for the pH experiment. This seemed reasonable because wetland soils go through flooding cycles, which seem to cause viral communities to become more mixed (more homogeneous over space) at different times of the year (ter Horst et al., 2023). However, here the wetland buffer chemistry biological replicates were even more spatially structured than the wetland PPBS buffer pH biological replicates (28.9% of vOTUs were shared between buffer chemistry replicates vs. 60.7% shared between pH replicates, **Figure 4E,F**), which was also reflected in significant differences in richness and DNA yields between biological replicates (**Figure S9B,E**). This suggests that, at least in the wetland soils tested here, buffer chemistry contributed to greater (but still small) differences in extractable viral community composition than did PPBS buffer pH, as the observed differences according to buffer chemistry were still significant across biological replicates even under more spatially heterogenous conditions in the field. It is important to note that the wetland soil site used for both the pH and buffer chemistry experiments was sampled 20 months apart, and this same soil at the two different time points exhibited very different physicochemical properties (**Figure S1C,D**). More specifically, the soil used in the pH experiment had significantly higher Ca^2+^ content (**Table S6**), which is thought to increase viral sorption (Sasidharan et al., 2016; Santos-Medellín et al., 2022), and thus could have introduced competing effects with the extraction efficiency of different buffer pHs. Specifically, pH would not have influenced differential sorption as substantially at greater Ca^2+^ concentrations, as divalent Ca^2+^ ions can bind viral particles more tightly than monovalent protons, especially at higher pHs (Kimura et al., 2008). Thus, while buffer chemistry seemingly played a more significant role than pH in the extraction of different wetland viral communities, we cannot rule out the possibility that these results were influenced by environmental differences in the soil used for each experiment.

## 4. DISCUSSION

Our understanding of soil viral ecology via viromics is based on the assumption that a representative consortium of viral particles can be efficiently extracted from diverse soils for downstream ecological analyses. However, viruses exhibit different sorption properties, according to soil and soil solution chemistry, which is why it has been suggested to tailor extraction buffers to different soil types, in order to extract as many viral particles as possible (Trubl et al., 2020). Here we showed that viral communities, measured via alpha- and beta-diversity, and viral particle abundances (by viromic DNA yield proxy), were overwhelmingly structured by sampling location only ∼1 m apart, and neither PPBS buffer pH (pHs 4.5, 5.5, 6.5, and 7.5) nor different buffer chemistries (PPBS, Carbonated Buffer, Glycine, and Saline Magnesium) changed observations of primary viral ecological patterns. This was consistent across the three soil types tested (forest, grassland, and wetland for PPBS buffer pH; buffer chemistry tests were only applied to the wetland), even though these soils differed significantly in physicochemical properties known to drive viral sorption efficacy (i.e., cation concentrations, organic matter content, (Jin and Flury, 2002)). Only the wetland soil biological replicates treated with different buffer chemistries were also significantly structured by extraction buffer chemistry, but this pattern was secondary to differences by sampling location. When controlling for spatial heterogeneity by homogenizing replicates and extracting from a pooled soil (i.e., technical replicates), the effects of buffer pH and chemistry became more apparent, but only for the grassland soil treated with PPBS at pH 4.5, and for the wetland soil treated with different extraction buffers. Taken together, extraction buffer as the primary driver of extracted viral community composition was only observed for homogenized soils, whereas spatial effects were the primary driver of viral community composition when a sampling distance of ∼1 meter was introduced. It is important to note that his sampling distance is much closer than distances between ‘replicates’ collected in other recent soil viral ecological studies (Santos-Medellín et al., 2021; Sorensen et al., 2021; ter Horst et al., 2021; Santos-Medellín et al., 2022; Durham et al., 2022; ter Horst et al., 2023; Hillary et al., 2022; Sorensen et al., 2023). These results demonstrate that viral ecological spatial patterns at very short distances are more prominent in structuring extractable viral communities than extraction buffer chemistry, and they suggest that viral communities collected across more heterogenous conditions (i.e., further spatial distances, different habitats, dynamic environmental conditions) would presumably be even less structured by buffer chemistry. For designing future field-based soil viral ecology studies via viromics, sampling locations, numbers of replicates, and distances among replicates appear to be much more important considerations than extraction buffer chemistry.

The pH of viromics extraction buffers is thought to be an important factor to consider for optimal viral particle extraction, due to changes in the interactions between viral surfaces and soil across different pHs (Powelson and Gerba, 1995; Williamson et al., 2003; Trubl et al., 2016). Generally, both soils and viral particles are negatively charged and repel each other, but with decreasing pH, different viral surface protein functional groups can become more protonated, causing some viral particles to become less negatively charged and subsequently sorbed to soil (Powelson and Gerba, 1995; Jin and Flury, 2002; Michen and Graule, 2010). It has therefore been suggested that the isoelectric point (IEP) of viral surface proteins (i.e., the pH at which a viral particle is neutrally charged) can be one of the best indicators of the sorption likelihood of a viral particle (Gerba et al., 1981; Dowd et al., 1998; Williamson et al., 2003; Trubl et al., 2016, 2020; Ghernaout and Elboughdiri, 2021), whereby a virus is expected to sorb to soil at pHs below its isoelectric point (Powelson and Gerba, 1995; Guan et al., 2003; Zhuang and Jin, 2003). Given this, we expected that extraction buffer pH should have selected for different pools of readily extractable viruses, which would have exhibited different protein compositions and related IEPs. However, across all PPBS extraction buffer pHs and soils tested (n = 72), only the grassland soil technical replicates at pH 4.5 (n = 3) exhibited different viral communities, meaning that changes to pH did not substantially affect the extractable viral particle pool. This was in part explained by the buffering capacity of the soils, where only the grassland soil reached all of the intended extraction buffer pHs. However, even within the grassland soil technical replicates, predicted surface protein IEPs were surprisingly not significantly different across extraction pHs. We believe this negative result is due to limitations in theoretical predictions of IEPs, the effectiveness of using viral surface proteins and IEPs for direct assessment of viral fate and transport, and the relative dearth of research on soil virus surface proteins overall. For example, some predicted IEPs can often be much higher than empirical IEPs, potentially due to nucleic acid charges or polynucleotide-binding regions being more influential to overall charge than just surface protein composition (Ghernaout and Elboughdiri, 2021; Heffron and Mayer, 2021), which may have muddled the trend expected in our IEP results. Further, electrostatic forces based on surface proteins are not the only factors at play when determining viral sorption, where viral particle size, hydrophobicity, conductivity of the soil solution, van der Waals forces, and many other factors can all largely affect the fate and transport of a virus (Powelson and Gerba, 1995; Jin and Flury, 2002; Ghernaout and Elboughdiri, 2021). Finally, there is still very little known about both soil virus surface proteins and the isoelectric points of these viruses. Many studies that have conducted empirical IEP research related to viral sorption in porous media have only done so on a handful of viruses (Dowd et al., 1998; Gitis et al., 2002; Chu et al., 2003; Guan et al., 2003; Davis et al., 2006), and our database only contained surface protein sequences from 1,493 unique viruses, whereas we uncovered 10,880 vOTUs (viral ‘species’), each presumably with at least one surface protein (e.g., a capsid protein). This highlights the need for more targeted research to expand the known soil virus protein composition and associated IEPs to better predict viral sorption in soils.

Different viromics extraction buffer chemistries used in the literature have demonstrated that extractable viral abundances can be largely influenced by the buffer used, which was also observed in the wetland soil tested here extracted under different buffer chemistries, but with opposite trends than previously described. For example, a previous study found that saline magnesium buffer was unable to recover most spiked-in phages (quantified via plaque-assay) when testing viromics buffer resuspension efficacy (Göller et al., 2020). However, in both the wetland technical and biological replicates tested here, saline magnesium extracted some of the highest concentrations of viral DNA (a proxy for viral particle abundance), as well as the second highest numbers of unique vOTUs in the biological replicates, after PPBS. Similarly, glycine buffer was found previously to recover the most viable viral particles inoculated into soil (quantified via plaque-assay, (Williamson et al., 2005)), whereas in our study, glycine extracted lower viral DNA yields compared to PPBS and saline magnesium, as well as the smallest number of unique vOTUs in the biological replicates. The different results from sequencing-based analyses in combination with viromic DNA yields as a proxy for abundance in this study compared to measurements focused on abundance alone (via microscopy or cultivation) in prior studies suggest that quantification and analysis methods likely alter conclusions regarding the efficiency of viral particle extraction by different viromics buffers. Overall, while the methods used in our study did reveal some differences in extractable wetland viral community composition and abundances based on extraction buffer used, most vOTUs were extracted by all buffers tested, and spatial distance was still the primary driver in the biological replicates. This means that, unlike what was observed in the studies discussed above, no particular extraction buffer was superior at viral particle extraction based on our quantification methods.

## 5. CONCLUSIONS

Here, we demonstrated that sampling location (samples collected ∼1 meter apart) was a more significant driver of extracted viral community composition than viromics extraction buffer chemistry (both different PPBS buffer pHs and buffer chemistries overall) in a forest, grassland, and wetland soil. When constraining spatial distances by homogenizing soil and extracting from technical replicates, the effects of PPBS buffer pH on viral community composition became more apparent, but only for the grassland soil. The grassland soil was unable to buffer changes in pH based on its edaphic properties in the technical replicates, presumably resulting in increased viral sorption to soil particles, but we could not confirm this trend by using predicted surface proteins and their isoelectric points, highlighting the need for more research on soil virus surface proteins to understand viral particle sorption dynamics. Given that the effect of PPBS buffer pH on soil viral community composition was only observed in one soil and only when spatial distances were constrained, results suggest that extraction buffer pH is not as important as previously thought.

However, different extraction buffer chemistries did significantly structure extractable viral communities in the wetland soil, for both technical and biological replicates, even though the biological replicates were more spatially heterogenous than those tested in the pH experiment. This implies that buffer chemistry could introduce substantial effects on recovered viral community composition from wetlands in more homogenous environmental conditions, which may confound commonly tested microbial ecology patterns (i.e., distance-decay).

However, spatial distance was still the prominent and most significant driver of viral community composition in the wetland biological replicates, which were collected over much smaller spatial scales (1 meter apart) than in other recent soil viral ecological studies. Thus, while the effects of extraction buffer (both different pHs and chemistries) are more prominent in highly homogenous soil environments (i.e., technical replicates, well-mixed wetlands), most natural habitats exhibit more heterogenous environmental conditions, and most viral ecological study samples are taken across larger spatial distances, both suggesting that extraction buffers do not substantially bias conclusions made from soil viral ecological studies.

## DATA AVAILABILITY

All raw sequences have been deposited in the NCBI Sequence Read Archive under the BioProject accession PRJNA1039020. All scripts are available at https://github.com/janefudyma/BufferChemistry.

## ACKNOWLEDGEMENTS

We thank Katherine Simpson-Johnson and Luke Salvato for statistical analysis help, and the University of California Natural Reserves site directors and staff, including Jacqueline Sones (Bodega Marine Reserve), Peter Steel (Angelo Coast Range Reserve), and Catherine Koehler (McLaughlin Reserve), for facilitating site access and providing logistical support. Funding for this work was provided by the U.S. Department of Energy (DOE), Office of Science, Office of Biological and Environmental Research (BER), Genomic Science Program, award number DE-SC0021198 (grant to JBE) and U.S. DOE-BER, Genomic Science Program (GSP), LLNL ‘Microbes Persist’ Soil Microbiome Scientific Focus Area SCW1632 (grant to JP-R, UC Davis subaward to JBE).

