## Supplementary Figures for "Exploring viral particle, soil, and extraction buffer physicochemical characteristics and their impacts on extractable viral communities"

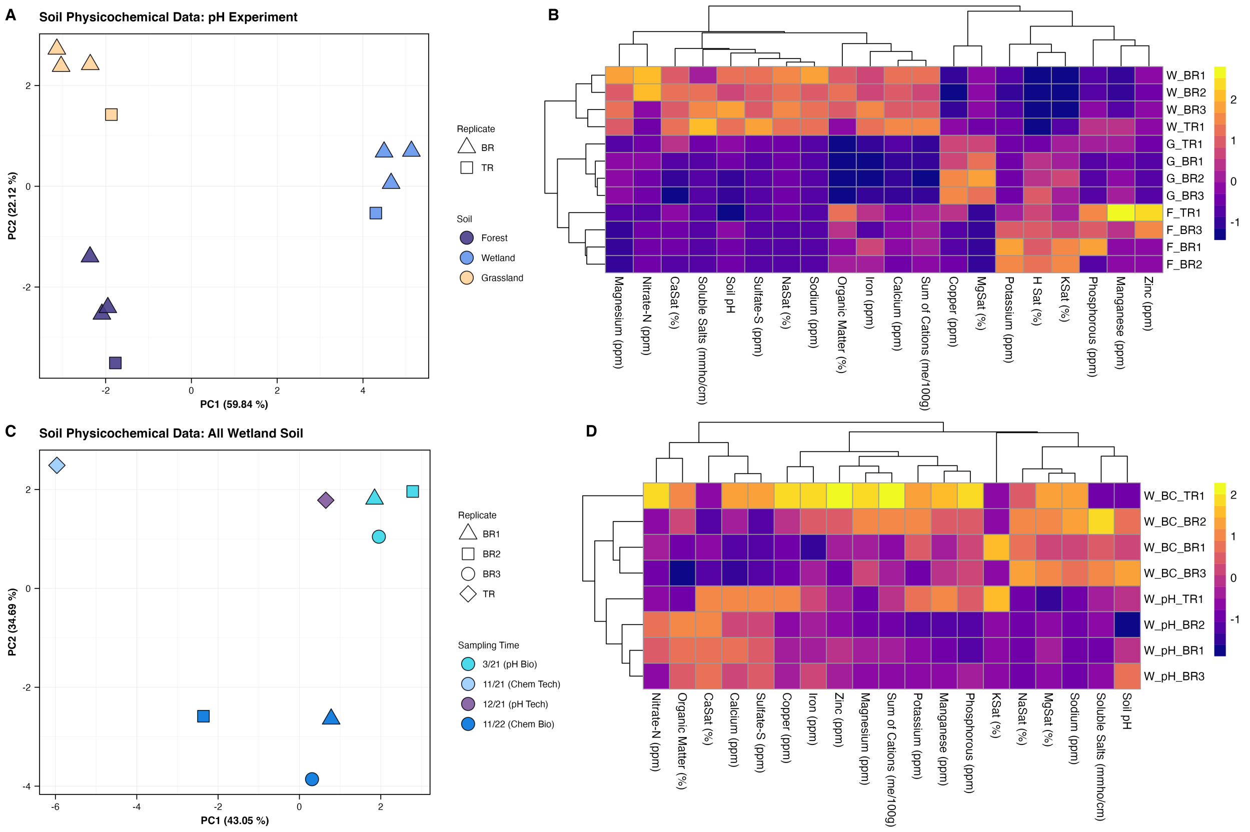


**Figure S1. A)** Principal components analysis (PCA) of z-score transformed Euclidean distances of soil physicochemical data (in panel B and Supplementary Table 1) for both the biological and technical replicates from the PPBS buffer pH experiments. Points reflect the physicochemical profile of one sample, colored by soil type. **B)** Hierarchal clustering of z-score transformed physicochemical data (columns) and samples according to physicochemical data (rows), visualized as a heatmap. Labels indicate the soil (W – wetland, G – grassland, F – forest) and the numbered replicate (BR – biological replicate, TR – technical replicate). **C)** Principal components analysis of z-score transformed Euclidean distances of wetland soil physicochemical data (in panel D and Supplementary Table 1) for the biological and technical replicates from both the PPBS buffer pH experiments and the buffer chemistry experiments. Points reflect the physicochemical profile of one sample, colored by soil type. **D)** Hierarchal clustering of z-score transformed physicochemical data (columns) and samples according to physicochemical data (rows), visualized as a heatmap. Labels indicate the soil (W – wetland), the type of experiment (pH – different PPBS buffer pHs, BC – different buffer chemistries), and the numbered replicate (BR – biological replicate, TR – technical replicate).

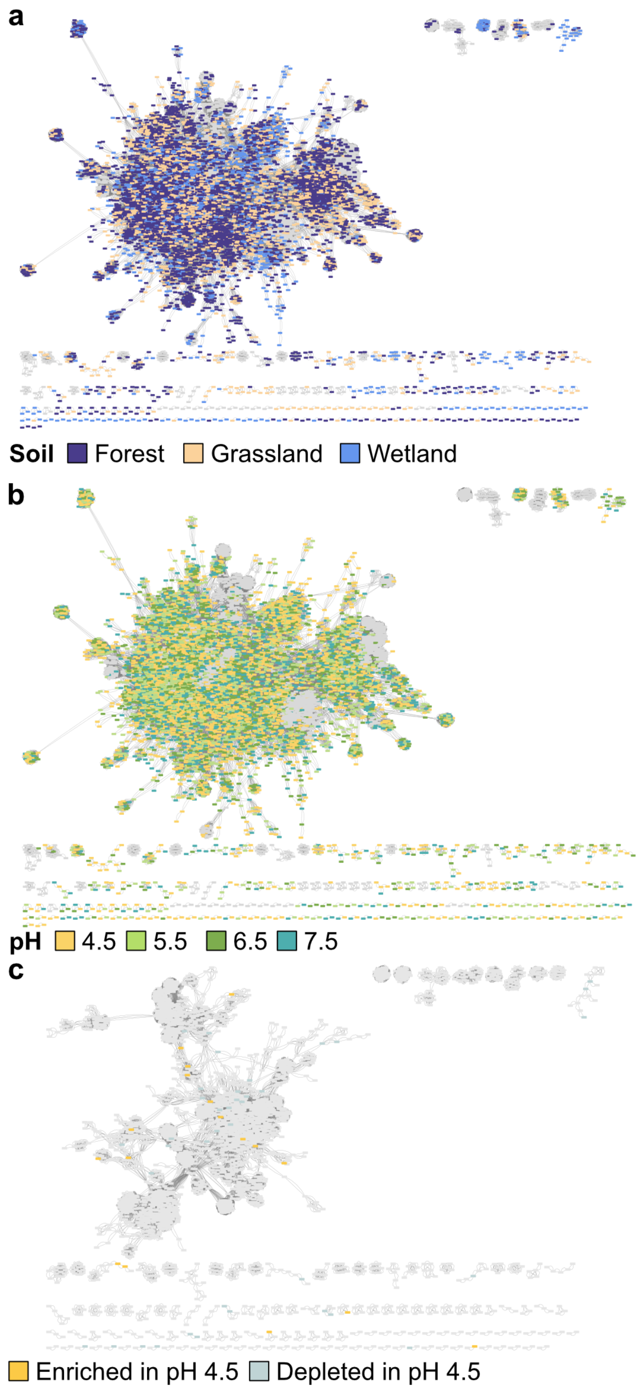

**Figure S2.** Gene-sharing network based on predicted proteins from recovered vOTUs in the PPBS buffer pH study. Nodes represent one vOTU and edges connect similar viral genomes according to predicted protein content. **A** & **B)** contain vOTUs from both biological and technical replicates recovered from all soils (n = 72, three soils, twelve replicates per soil, two replicate types), and each vOTU is colored based on **A)** the soil from which the vOTU assembled or **B)** buffer pH from which the vOTU assembled. **C)** contains vOTUs from only the grassland technical replicates, colored by significantly differentially abundant vOTUs in the pH 4.5 extraction (i.e., either enriched or depleted in pH 4.5 as compared to 5.5, 6.5 and 7.5).

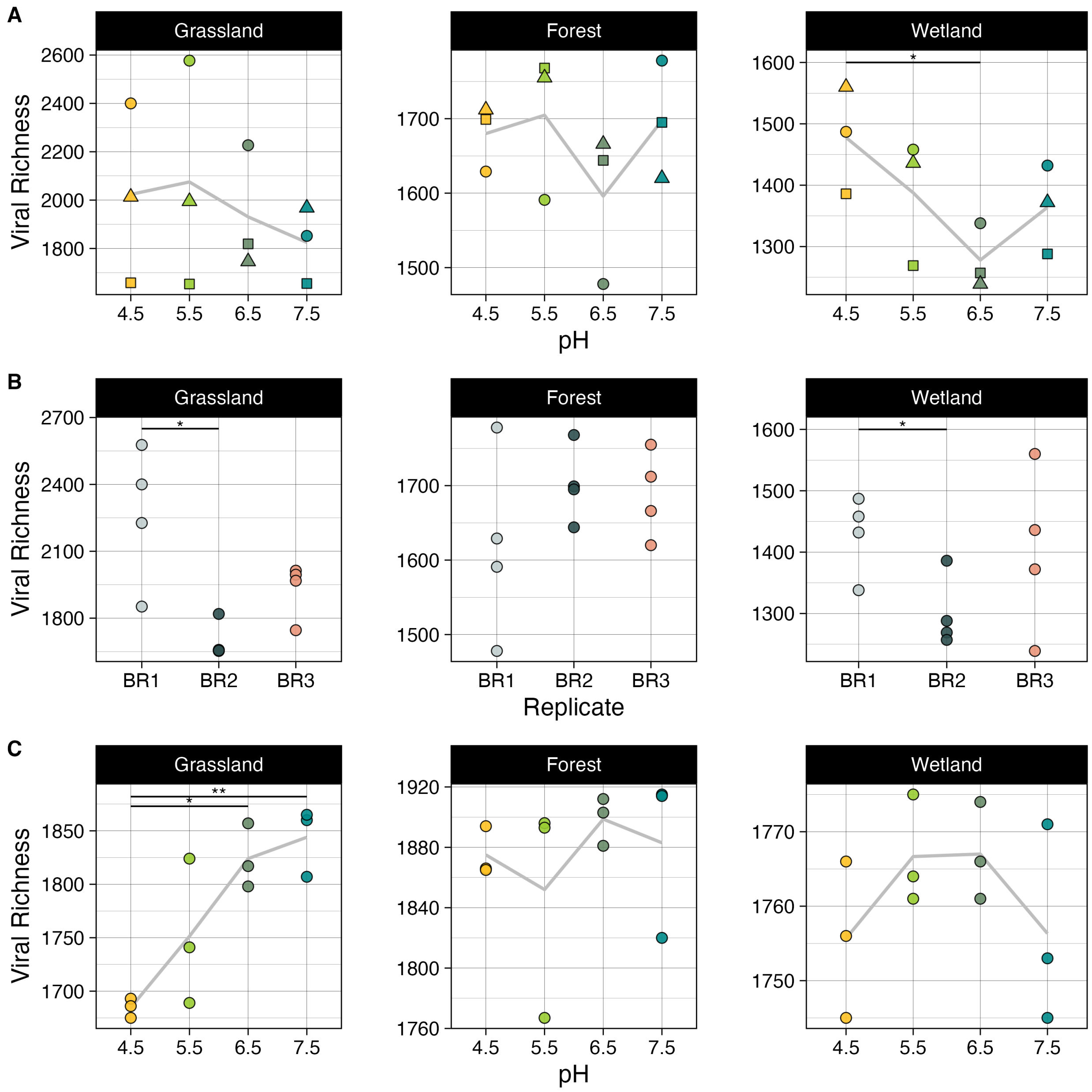


**Figure S3.** Viral community richness (total number of vOTUs) for the two PPBS buffer pH experiments faceted by soil type, reflecting differences in richness among **A)** biological replicates across PPBS buffer pHs, with shapes reflecting replicate replicate (circles – biological replicate 1, squares – biological replicate 2, triangles – biological replicate 3), **B)** biological replicates, and **C)** technical replicates across PPBS buffer pHs. Trend lines in **A&C** follow the mean viral richness across buffer pHs. Significance between groups is marked by connecting lines with significance stars (*, p <0.05; ** p <0.01), based on analysis of variance (ANOVA) and Tukey post hoc test.

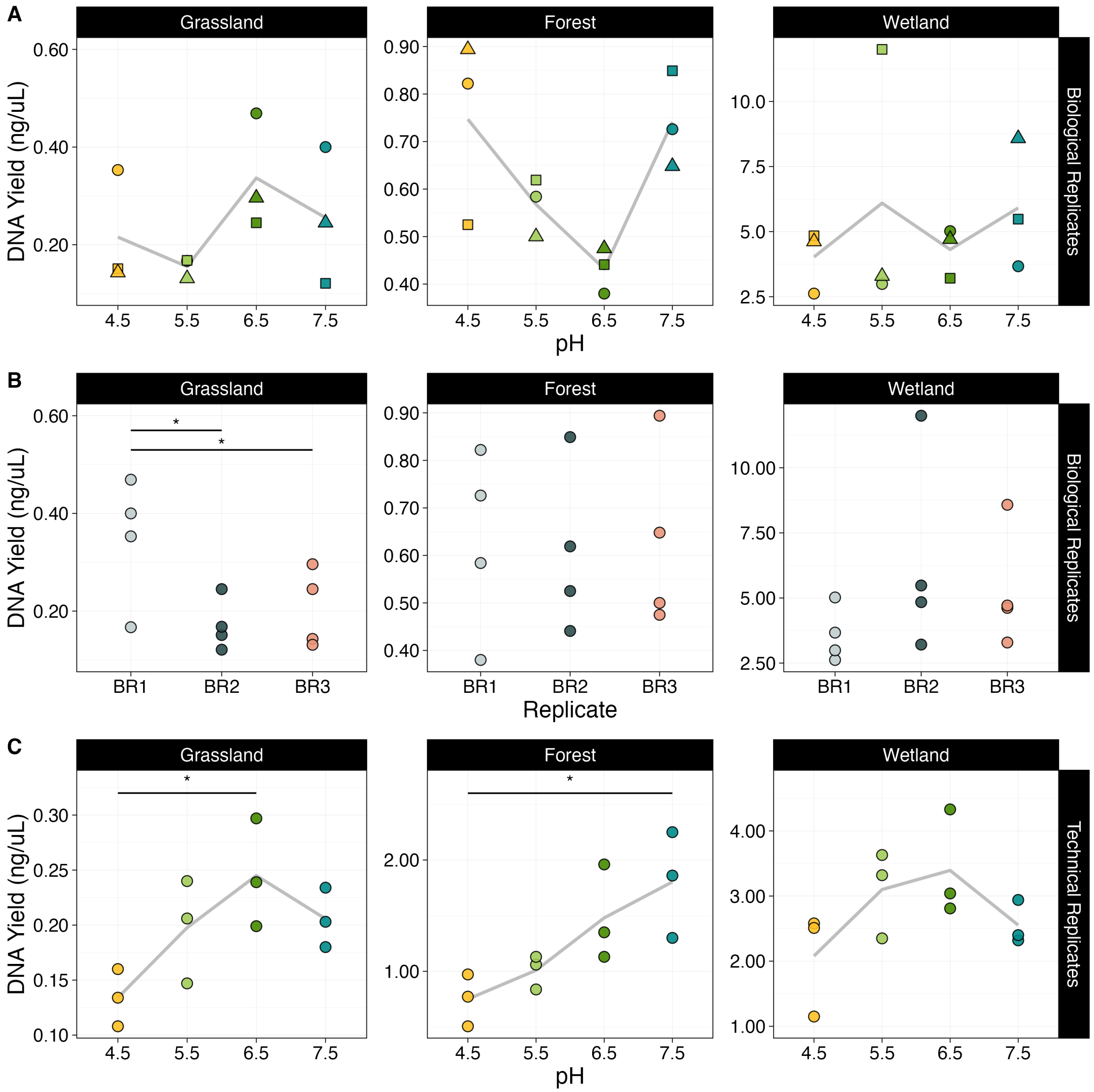


**Figure S4.** Viromic DNA yields (a relative proxy for viral particle abundances) for the two PPBS buffer pH experiments faceted by soil type, showing trends across **A)** buffer pH for biological replicates, with shapes reflecting replicate (circles – biological replicate 1, squares – biological replicate 2, triangles – biological replicate 3), **B)** replicates for biological replicates, and **C)** buffer pH for technical replicates. Trend lines in **A&C** follow the mean DNA yield concentrations across buffer pHs. Significance between groups is marked by connecting lines with significance stars (*, p <0.05), based on analysis of variance (ANOVA) and Tukey post hoc test.

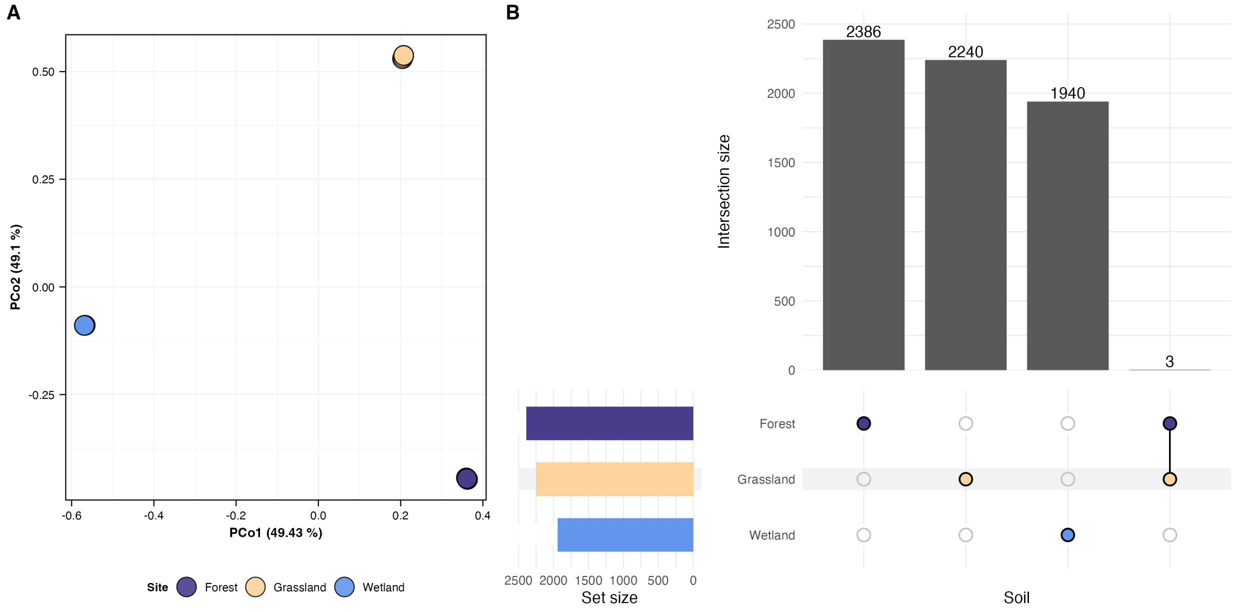


**Figure S5. A)** Principal coordinates analysis (PCoA) based on Bray-Curtis dissimilarities of vOTU abundances (coverage depths from read mapping) from three soils, with 12 viromes per soil (three technical replicate samples collected ~1 meter apart, homogenized, and subsampled from the homogenate, then treated with protein-supplemented phosphate-buffered saline solution (PPBS) at four different pHs: 4.5, 5.5, 6.5, and 7.5). Each point is one virome, representing one technical replicate and pH treatment combination, colored by soil type. **B)** UpSet plot of shared vOTUs in each soil type, based on presence-absence data derived from read mapping to vOTUs. Colored dots indicate the soil type(s) in which a given set of vOTUs was detected, and connecting lines between dots indicate vOTUs shared between soils. Data for each soil is derived from 12 samples (three technical replicates and four PPBS buffer pHs), and vOTUs detected in any sample from a given category were counted in this analysis.

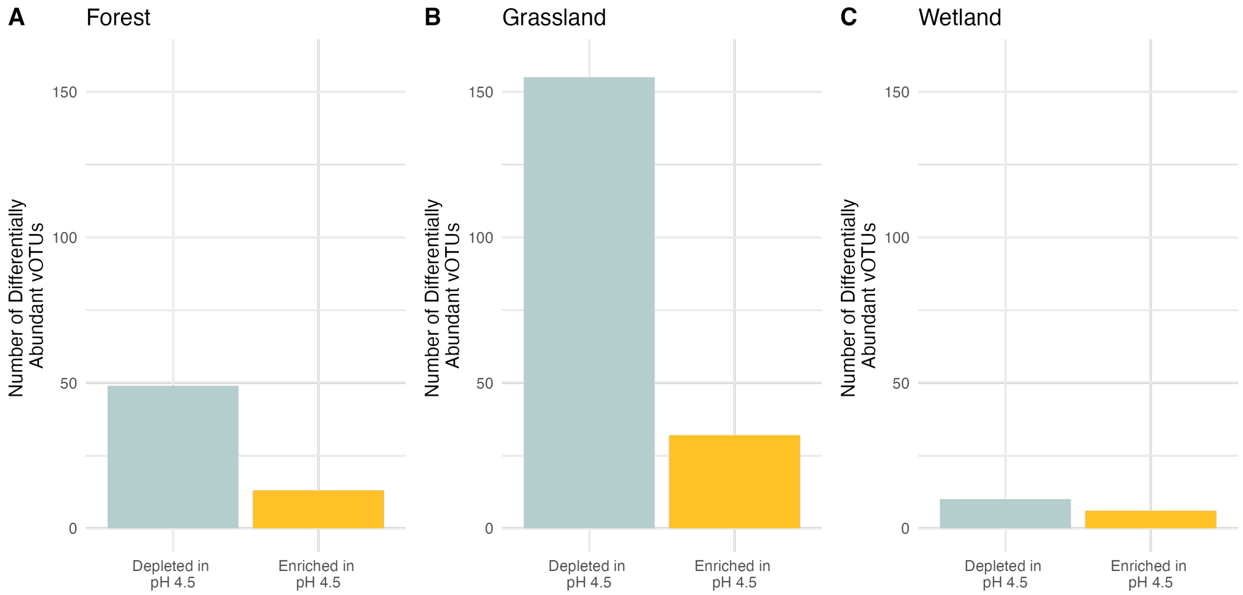


**Figure S6.** Significant differentially abundant vOTUs from technical replicates, either enriched or depleted in pH 4.5 PPBS buffer, as compared to pH 5.5, 6.5, and 7.5, in the **A)** Forest **B)** grassland, and **C)** wetland soils. P-values to determine differential abundance significance of vOTUs between groups were first generated from pairwise Wald tests and subsequently adjusted using the Bonferroni method to account for all comparisons.

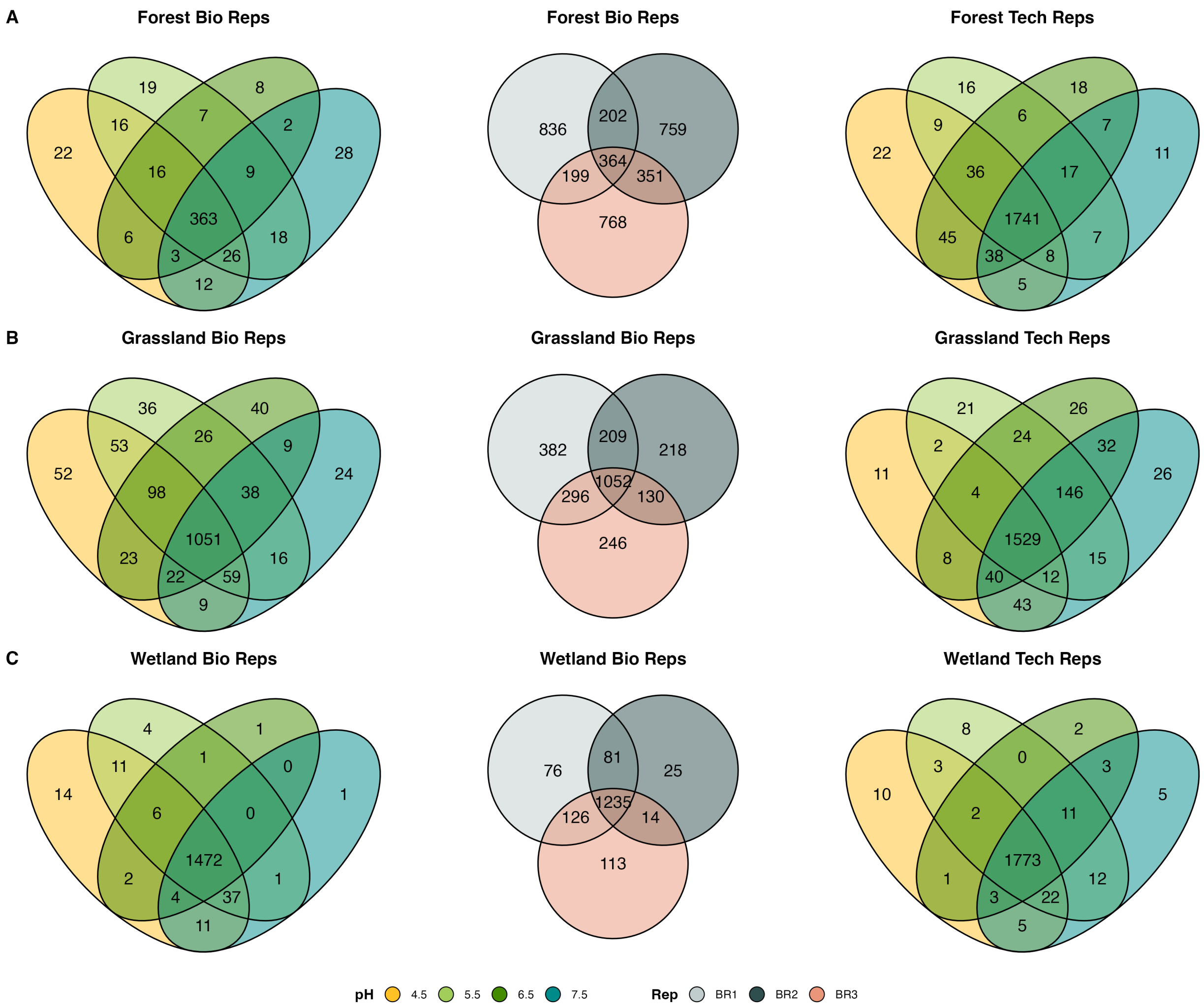


**Figure S7.** Venn diagrams of vOTU detection patterns from the PPBS buffer pH experiment (based on presence-absence data derived from read mapping to vOTUs); vOTUs were only considered for this analysis if they were detected in all technical or biological replicates for a given buffer treatment, or in all treatments for a given replicate. vOTUs from **A)** the forest soil, detected within or across different PPBS buffer pHs or biological replicates, **B)** the grassland soil, detected within or across different PPBS buffer pHs or biological replicates, and **F)** the wetland soil, detected within or across different PPBS buffer pHs or biological replicates.

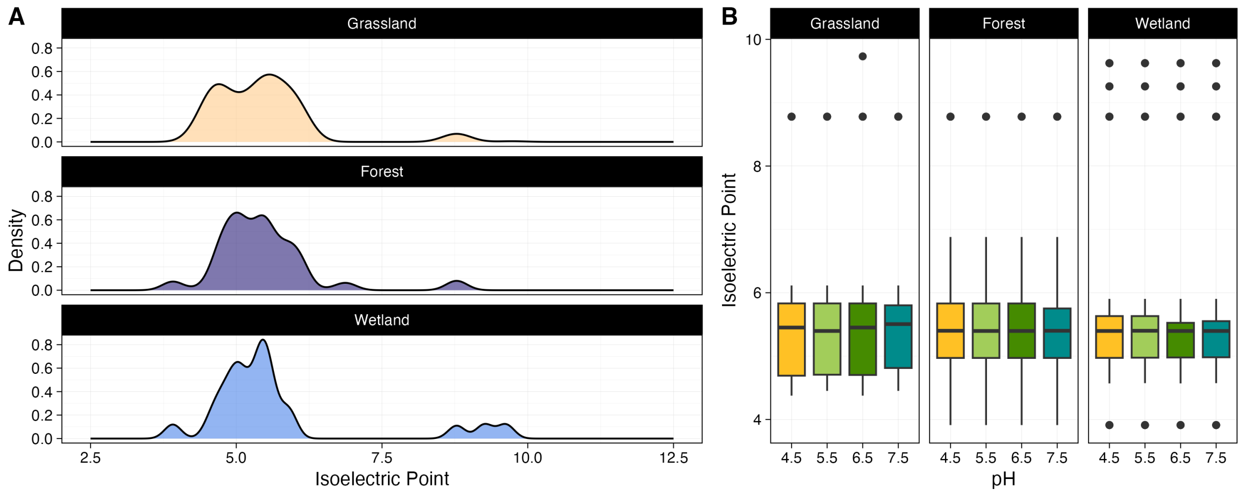


**Figure S8. A)** Density plots reflecting distributions of bioinformatically calculated isoelectric points (i.e., the pH at which a protein is neutrally charged) from predicted viral surface proteins of vOTUs recovered from all soil technical replicates from the PPBS buffer pH experiment, faceted by soil type. All recovered protein isoelectric points were used in this analysis, regardless of the buffer pH in which they were recovered. **B)** Box plots of surface protein isoelectric points across different buffer pHs in the technical replicates, faceted by soil type. Each box represents all isoelectric points predicted for a protein recovered at a given pH, lines indicate medians, boxes indicate the 25$^{th}$ and 75$^{th}$ percentiles, and whiskers represent the ±1.5 interquartile range.

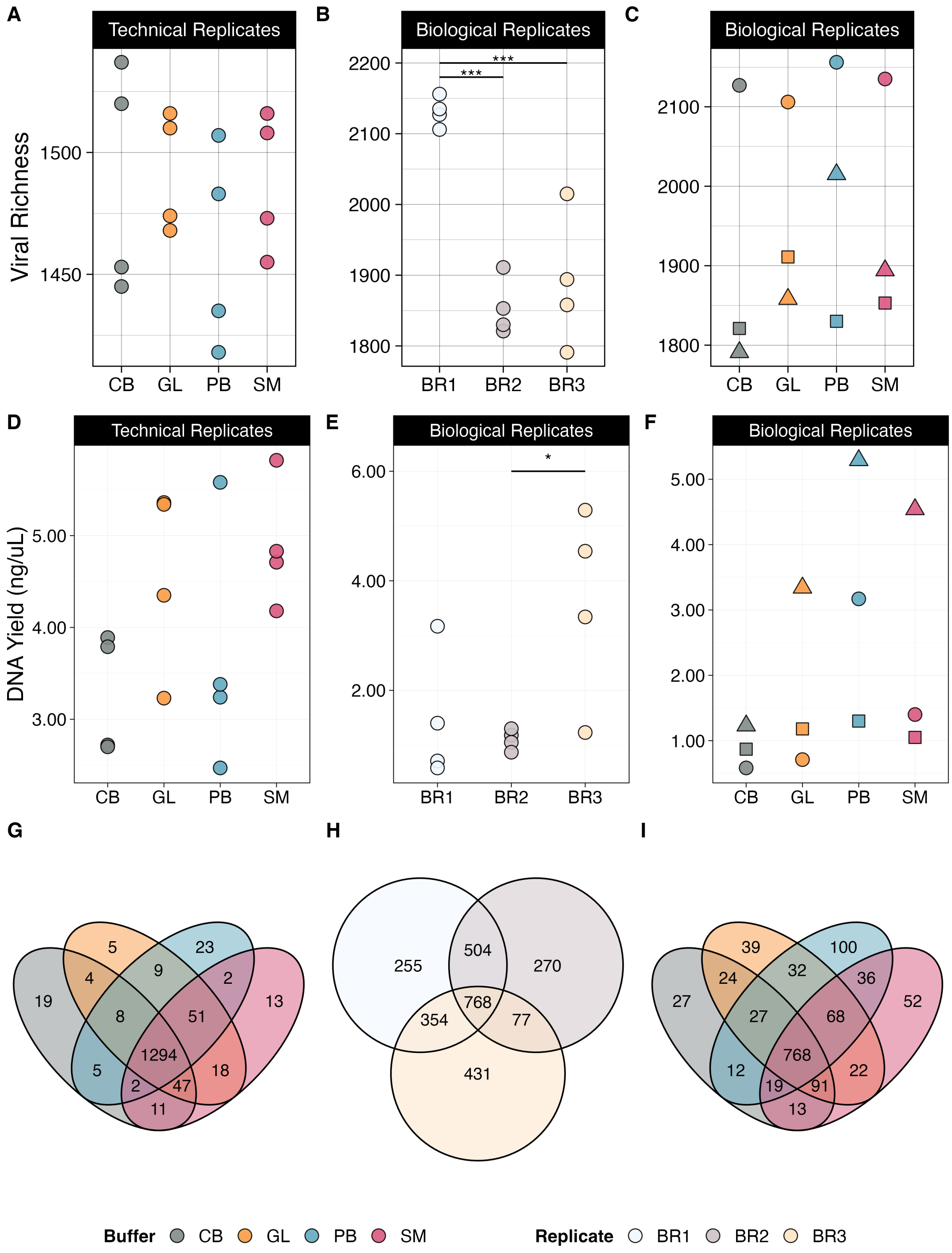


**Figure S9. A-C)** Viral community richness (total number of vOTUs) measured in the wetland soil for **A)** technical replicates across different buffer chemistries [CB – carbonated buffer, GL – glycine, PB – protein-enhanced phosphate buffered saline (PPBS, as in earlier tests, here pH 6.5), SM – saline magnesium], **B)** biological replicates by replicate, and **C)** biological replicates across different buffer chemistries. **D-F)** Viromic DNA yields (a relative proxy for viral particle abundances) from the wetland soil buffer chemistry experiments (both technical and biological replicates), showing trends across **D)** different buffers for technical replicates, **E)** replicates for biological replicates, and **F)** different buffers for biological replicates. In **C&F**, shapes reflect replicate number (circles – biological replicate 1, squares – biological replicate 2, triangles – biological replicate 3). In **A-F**, significance between groups is marked by connecting lines with significance stars (*, $p < 0.05$; ***, $p < 0.001$), based on analysis of deviance and Tukey post hoc test. **G-I)** Venn diagrams of wetland vOTU detection patterns (based on presence-absence data derived from read mapping) within or across **G)** all four buffer chemistries in the technical replicates, **H)** all replicates in the biological replicates and **I)** all four buffer chemistries in the biological replicates. vOTUs were only considered for this analysis if they were found in all biological replicates for a given buffer treatment, or all treatments for a given biological replicate.
